# Auxin-induced expression divergence between Arabidopsis species likely originates within the TIR1/AFB-AUX/IAA-ARF module

**DOI:** 10.1101/038422

**Authors:** Jana Trenner, Yvonne Poeschl, Jan Grau, Andreas Gogol-Döring, Marcel Quint, Carolin Delker

## Abstract

TIR1/AFB, AUX/IAA, and ARF proteins show interspecies expression variation correlating with variation in downstream responses which indicates a source for natural variation within this conserved signaling module.

**Abstract:** Auxin is an essential regulator of plant growth and development and auxin signaling components are conserved among land plants. Yet, a remarkable degree of natural variation in physiological and transcriptional auxin responses has been described among *Arabidopsis thaliana* accessions. As intra-species comparisons offer only limited genetic variation, we here inspect the variation of auxin responses between *A. thaliana* and *A. lyrata.* This approach allowed the identification of conserved auxin response genes including novel genes with potential relevance for auxin biology. Furthermore, promoter divergences were analyzed for putative sources of variation. *De novo* motif discovery identified novel and variants of known elements with potential relevance for auxin responses, emphasizing the complex, and yet elusive, code of element combinations accounting for the diversity in transcriptional auxin responses. Furthermore, network analysis revealed correlations of inter-species differences in the expression of *AUX/IAA* gene clusters and classic auxin-related genes. We conclude that variation in general transcriptional and physiological auxin responses may originate substantially from functional or transcriptional variations in the TIR1/AFB, AUX/IAA, and ARF signaling network. In that respect, *AUX/IAA* gene expression divergence potentially reflects differences in the manner in which different species transduce identical auxin signals into gene expression responses.

## Introduction

Auxin’s capacity to regulate the essential cellular processes of division, elongation and differentiation integrates it in the regulation of virtually all developmental and physiological plant processes. On a molecular level, auxin responses involve extensive and rapid changes in the transcriptome (Paponov *et al.*, 2008). This response depends on a signaling pathway which is constituted by three main signaling components: (i) TRANSPORT INHIBITOR RESPONSE1/AUXIN SIGNALING F-BOX 1-5 (TIR1/AFBs) auxin-co-receptors, (ii) AUXIN/INDOLE-3-ACETIC ACID (AUX/IAA) family of auxin co-receptors/transcriptional repressors, and (iii) the AUXIN RESPONSE FACTOR (ARF) family of transcription factors (Quint and Gray, 2006).

ARFs induce or repress the expression of genes by binding to auxin-responsive elements (AuxRE) in the respective promoter regions (Guilfoyle *et al.*, 1998; Ulmasov *et al.*, 1999). When auxin levels are low, AUX/IAAs prevent ARF regulatory action on auxin-responsive genes (Weijers *et al.*, 2005; Szemenyei *et al.*, 2008). The presence of auxin is sensed by a co-receptor complex formed by the cooperative binding of auxin by the TIR1/AFB F-box subunit of an SCF-type E3 ligase and an AUX/IAA protein (Dharmasiri *et al.*, 2005; Kepinski and Leyser, 2005; Calderón Villalobos *et al.*, 2012). This binding results in the polyubiquitylation of the AUX/IAAs by the SCF^TIR1/AFB^ complex (Maraschin *et al.*, 2009). The subsequent proteasomal degradation of the tagged AUX/IAAs causes a de-repression of ARF transcription factors, which are then released to initiate transcriptional changes (Ramos *et al.*, 2001; Zenser *et al.*, 2001). The three key signaling elements of TIR1/AFBs, AUX/IAAs, and ARFs are encoded by gene families of six, 29 and 23 members, respectively (Chapman and Estelle, 2009). Numerous possibilities of combinations among the individual gene family members with putatively different signaling capacities could ultimately be responsible for the wide range of auxin signaling outputs observed throughout plant growth and development (Calderon Villalobos *et al.*, 2012; Salehin *et al.*, 2015).

The auxin signaling pathway is conserved among land plants as individual core components are present already in the liverwort *Marchantia polymorpha* (Kato *et al.*, 2015). With the universal impact of auxin on plant growth and development, an open question in auxin biology remains whether auxin signaling and response contribute to adaptive processes to local environmental conditions and challenges. First data indicating that the read-out of an auxin stimulus can be highly variable were obtained by the analysis of natural variation of auxin responses among different natural accessions of *A. thaliana* (Delker *et al.*, 2010). Variation in transcriptomes and co-expression networks of signaling elements gave rise to the hypothesis that altered equilibria of individual signaling components might contribute to the variation observed on the general transcriptome and ultimately on the physiological level (Delker *et al.*, 2010).

Here, we performed a cross-species analysis of auxin responses in the two closely related species *A. thaliana* and *A. lyrata* in a comparative transcriptomics approach. The increased genetic variation between the two *Arabidopsis* species compared to the variation among different accessions allowed (i) the identification of conserved auxin response genes, (ii) the identification of *cis*-regulatory elements that might contribute to auxin responses, (iii) to assess whether the hypothesized variation in early auxin signaling gene expression as a source for downstream variation could be verified in a system with higher genetic variation, and (iv) the comparison of inter- and intra-species variation of *AUX/IAA* gene clusters and downstream variation.

## Materials and Methods

### Plant material and growth conditions

*A. thaliana* Col-0 (N1092) and *A. lyrata* (N22696) were obtained from the Nottingham Arabidopsis Stock Centre. Seeds were surface-sterilized and stratified 3 d at 4 °C before sowing. Seedlings were grown on solid or in liquid *Arabidopsis thaliana* solution (ATS) nutrient medium (Lincoln *et al.*, 1990). For growth assays, seedlings were cultivated on vertical ATS for 3 d (IAA), 4 d (TIHE), or 5 d (2,4-D and NAA) before transfer to plates supplemented with IAA, 2,4-D, or NAA at the indicated concentrations or before transfer of plates to 28 °C (TIHE). Root lengths were quantified after an additional 5 d (IAA) or 3 d (2,4-D and NAA), hypocotyl growth was quantified after additional 4 d at 28 °C. All experiments were performed in long-day conditions (16 h light/8 h dark) and a fluence rate of ~ 230 μmol m^-2^ sec^-1^ (root growth assays) or 30 μmol m^-2^ sec^-1^ (TIHE). Relative root and hypocotyl lengths of hormone- and temperature-treated seedlings, respectively, were determined as percent in relation to the median value of 20 °C grown plants. Statistical analyses (1- and 2-way ANOVAs) were performed on the untransformed raw data. For expression studies and [^3^H]-IAA uptake assays, seeds were cultivated in liquid ATS under continuous illumination to minimize potential circadian effects. For expression analyses, ATS was supplemented with 1 μM IAA for 0, 1 h, and 3 h after seven days. Yellow long-pass filters were applied in all IAA treatment experiments to prevent photodegradation of IAA.

### [^3^H]-IAA uptake assay

Three biological replicates of seven days-old seedlings were treated with 2 nM of [^3^H]-IAA (Hartmann Analytic, Germany) per mg seedling fresh weight in liquid ATS for 1 h. Samples were subsequently washed with liquid ATS ten times before quantification via scintillation count.

### RNA extraction and microarray hybridization

RNA was extracted from three biological samples of seven days-old whole seedlings using the RNeasy Plant Mini Kit (Qiagen). RNA samples were further processed and hybridized to the ATH1-121501 microarray by the Nottingham Arabidopsis Stock Centre’s microarray hybridization service.

### Probe masking, data normalization and data processing

The raw data generated by NASC was pre-processed and corrected according to (Poeschl *et al.*, 2013) including the proposed polynomal correction of probe intensities. The data matrix contained the expression values for 16315 genes at three time points (with three biological replicates each) for both species. Significant changes in auxin-induced expression were determined by a modified t-test (Opgen-Rhein and Strimmer, 2007). P-values were Benjamini-Hochberg-corrected for multiple testing and genes significantly (fdr < 5%) changed by a factor of two or more (|log2 fold change| > 1) where considered to be differentially expressed.

### Modified Pearson correlation

To incorporate the information on variation among the biological replicate measurements at each time point in the correlation analyses, a modified Pearson correlation coefficient *(mod.r)* was introduced. 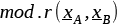 of the expression profiles for two genes A and B was computed by dividing the covariance of the mean expression profiles 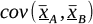 by the product of the standard deviations of the expression profiles 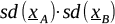, which is given by the formula:

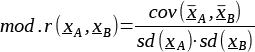

The mean expression profiles 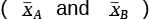 consist of one value per time point, which represent the means of the respective replicates.

### Cluster Analysis

A total of N=9091 genes were selected based on a coefficient of variation (cv) in expression profiles of cv > 0.05. A hierarchical clustering with average linkage was performed on N expression profiles using *1-mod.r* as distance measure. Each expression profile consists of 18 measurements representing the three biological replicates of three time points and two species. The resulting dendrogramm was cut level 0.1 *(mod.r* = 0.9) and resulting clusters were subsequently filtered by the following parameters: Clusters needed to contain at least five genes of which 70% showed a significant difference in species, time point and interaction as assessed by two-way ANOVAs which resulted in 14 clusters containing 337 genes in total.

### Promoter Analysis

Promoter sequences for *A. thaliana* and *A. lyrata* were extracted using the annotation provided by Phytozome v7.0 (http://www.phytozome.com). A promoter sequence was defined as 500 bp upstream the transcription start site to 100 bp downstream the transcription start site, or to the start codon, whichever came first.

### Extraction and assignment of known cis-elements

Extracted promoter sequences were analyzed for the presence of a set of annotated *cis*-elements and their reverse complements from http://arabidopsis.med.ohio-state.edu/AtcisDB/bindingsites.html (last accessed 2014/02/03) extended by a set of 10 *cis*-elements described in literature to be involved in auxin response/signaling (Tab. S2). Motifs shorter than six bases were excluded from the analysis. The sequences of the motifs were used as regular expressions to compute their occurrences in the promoter sequences.

### Determination of promoter and expression divergence

Similarities of promoter sequences of an orthologous gene pair was assessed by determining the occurrence of each possible 8-mer in each of the two promoter sequences and computing the Pearson correlation coefficient of the two vectors of k-mer counts (*kmer.r*) as proposed in (Vinga and Almeida, 2003). Promoter divergence was assessed as 1-*kmer.r* and expression divergence was determined as 1-*mod.r.*

### De-novo identification of putative cis-elements

Dimont (Grau *et al.*, 2013) was used for identification of putative novel *cis*-elements with slight modifications from the published procedure which are comprehensively described in the Supplemental Methods Section.

### Co-expression Analysis using Profile Interaction Finder (PIF)

The *Profile Interaction Finder* algorithm (PIF; Poeschl *et al.*, 2014)) was applied in its second mode using eight input profiles of the individual mean expression profiles of the eight *AUX/IAA* gene clusters. We applied the PIF to the set of genes showing a cv > 0.05 to prevent false-positive correlation based on noise. Parameters and thresholds for the identification of positively or negatively correlated genes were set to a |PIF-correlation| of > 0.7, neighbor number k = 1 and a 75 % bootstrap occurrence (n=1000). The PIF was additionally applied to nine input profiles of an intra-species *AUX/IAA* gene clusters analysis using the same conditions as for the inter-species analysis described prior.

### Statistical and computational analyses

Analyses were performed using the software R (R Core Team, 2015) with implementation of the following packages: beeswarm (Eklund, 2015), gplots (Warnes *et al.*, 2014), st (Opgen-Rhein and Strimmer, 2007), multtest (Pollard *et al.*, 2005).

### Accession numbers

The cross-species hybridization microarray data analyzed in this article are publicly available at http://data.iplantcollaborative.org/quickshare/8e9b2f0212c8a1bc/Exp579.zip.

## Results and Discussion

We inspected inter-species variation of auxin responses between *A. thaliana* and *A. lyrata* taking advantage of the close relation of the two *Arabidopsis* species whith extensive synteny despite considerable genetic variation, for example in total genome size (Hu *et al.*, 2011). The aim was to combine physiological, transcriptomic and genomic information to assess the extent of inter-species variation in auxin responses on several levels and to identify genes with conserved transcriptional responses. Furthermore, we wanted to exploit the genetic variation among the two sister species to gain further insights into the molecular mechanisms that contribute to naturally occurring variation in auxin responses which might ultimately reflect consequences of adaptation processes.

### Physiological auxin responses

To assess whether *A. thaliana* and *A. lyrata* show differences in physiological auxin responses, we used classic auxin response assays that focus on the quantitative reaction of seedling growth to exogenously applied auxin or to a temperature-induced increase of endogenous auxin levels. We performed several of these assays, testing the response to the naturally prevalent auxin indole-3-acetic acid (IAA) as well as several synthetic auxins, to assess the extent of natural inter-species variation between *A. thaliana* and *A. lyrata*.

In terms of relative growth effects, a high diversity in responses to natural and synthetic auxins was observed (Fig. 1A-D). While *A. thaliana* is less sensitive with respect to IAA-induced root growth inhibition (Fig. 1A), a higher sensitivity in temperature-induced hypocotyl elongation (TIHE) was observed (Fig. 1D). *A. thaliana’s* response to the synthetic auxin 2,4-Dichlorophenoxyacetic acid (2,4-D) was significantly stronger than the response of *A. lyrata* (Fig. 1B). In contrast, 1-Naphthaleneacetic acid (1-NAA)-induced root growth inhibition was almost similar in both species (Fig. 1C). Overall, the extent of variation in auxin responses between *A. thaliana* and *A. lyrata* seems to be highly dependent on the specific auxinic compound and the analyzed organ. The compound- and tissue-specificity might indicate differential sources for the observed response differences putatively involving any or all aspects of auxin biology ranging from biosynthesis (in case of TIHE) to transport, sensing, signal transduction and/or metabolism.

**Fig. 1.**
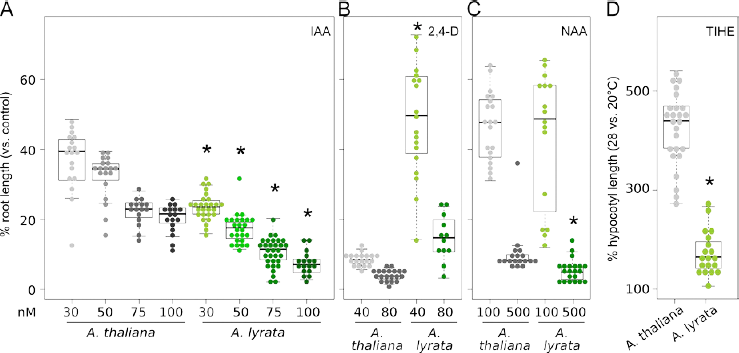
Physiological auxin responses of *A. thaliana* and *A. lyrata*. Relative root length (treated vs. control) of seedlings grown on different concentrations of (A) IAA, (B) 2,4-D, or (C) NAA. Three (A) or five (B,C) days-old seedlings were transferred to hormone-containing medium and grown for additional five (A) or three (B,C) days. (D) Relative hypocotyl length (28 °C/20 °C) of eight days-old seedlings. Box plots show medians (horizontal bar), interquartile ranges (IQR, boxes), and data ranges (whiskers) excluding outliers (defined as > 1.5 x IQR). Individual data points are superimposed as beeswarm plots. Asterisks denote significant differences between treatment responses of *A. thaliana* and *A. lyrata* as assessed by two-way ANOVA (i.e. genotype x treatment effect, P < 0.05) of the absolute data presented in Fig. S1.

### Microarray-based transcriptional profiling of auxin responses

For *A. thaliana,* natural variation among different accessions was observed on physiological as well as on transcriptional levels (Delker *et al.*, 2010). We thus conducted a microarray-based analysis of transcriptional auxin responses comparing *A. thaliana* and *A. lyrata* using a cross-species hybridization approach. We chose the same experimental set up as in previously reported transcriptome profiling studies in *Arabidopsis thaliana*

(Nemhauser *et al.*, 2006; Delker *et al.*, 2010) to maximize comparability among available data. In brief, seven days-old seedlings grown in liquid culture were treated with 1 μM IAA for one and three hours, respectively. Isolated RNAs from treated and control (untreated) seedlings were subsequently processed and hybridized to the Affymetrix ATH1 microarray. To exclude potential effects of differential auxin uptake on the transcriptional read-out, we quantified the amount of radio-labeled auxin in seven days-old seedlings exposed to [^3^H]-IAA for one hour (Fig. 2A, Supplemental Fig. S1G). The lack of statistically significant differences in [^3^H]-IAA levels indicated similar IAA uptake capacities in *A. thaliana* and *A. lyrata*. To further omit putative effects of differential internal transport we applied IAA in a concentration (1 μM) that is high enough to ensure saturation. The hybridization of a non-intended species to a species-specific microarray requires a probe-masking procedure in the processing of the expression data to avoid false-positive and false-negative results caused by mis-hybridization of probes due to sequence variations between the two species. Here, a sequence-based masking approach was applied that allows for one mismatch per probe and retained only those genes that are represented by at least three probes per probe set and uniquely hybridize to orthologous genes in *A. thaliana* and *A. lyrata* (Poeschl *et al.*, 2013). As a result of the masking procedure, 16315 genes were retained for expression comparisons between *A. thaliana* and *A. lyrata*. To correct for putative effects of one tolerated mismatch per probe on the expression level we implemented a fourth-degree polynomial correction option in the RMA-normalizing procedure as suggested by Poeschl et al. (2013). After normalization we inspected the expression levels of various constitutively expressed genes designated as superior expression reference genes in *A. thaliana* (Czechowski *et al.*, 2005) This subset of genes showed similar transcription profiles as well as largely similar expression levels in both *Arabidopsis* species indicating the comparability of the two data sets (Fig. S2).

**Fig. 2.**
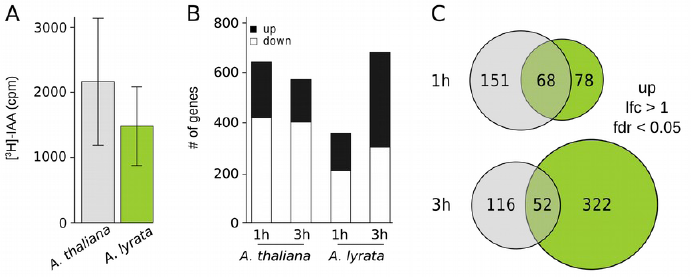
Quantification of [^3^H]-IAA uptake and ATH1-based assessment of auxin-induced transcriptome changes. (A) Seven days-old seedlings were treated with 2 ng [^3^H]-IAA per mg seedling fresh weight for 1 h in liquid ATS medium. Scintillation counts were recorded after removal of radiolabeled IAA and ten subsequent wash steps with liquid ATS. Bar plots show mean [^3^H]-IAA levels of three biological replicates and error bars denote SEM. No significant differences were detected by a two-sided t-test (P < 0.05). Results of a second independent experiment are shown in Supplemental Fig. S1G. (B) Stacked bars show the number of up- and down-regulated genes with an auxin-induced significant (fdr ≤ 0.05) change in expression level in black and white, respectively. (C) Venn diagrams illustrate the number of genes commonly or specifically up-regulated in *A. thaliana* (gray) and *A. lyrata* (green) after 1 h and 3 h of auxin treatment (lfc = log_2_ fold change, fdr = false discovery rate). The complete list of genes is presented as Supplemental Data File 1.

To analyze auxin-induced transcriptome changes, differentially expressed genes in both species were identified based on a significant (fdr < 0.05) change in expression with a |log2 fold change| > 1. Several hundred genes were differentially regulated in response to auxin in both species (Fig. 2B). Considerably more genes were differentially regulated in *A. thaliana* in response to one hour of auxin treatment than in *A. lyrata,* whereas after three hours more genes were responsive in *A. lyrata*. Overall, the number of down-regulated genes was relatively high in comparison to other auxin response transcriptome analyses (Paponov *et al.*, 2008; Delker *et al.*, 2010) In accordance with previous studies, we focused primarily on differentially up-regulated genes in the subsequent analyses. To further assess whether a time-delay in response may be a factor in creating the diverse response pattern we performed cross-comparison among the up-regulated genes after 1 h in *A. thaliana* and 3 h in *A. lyrata* (and vice versa). In case of a delayed response a considerable increase in the overlapping genes should occur in one of the cross-comparisons. However, as this is not the case (Supplemental Fig. S1H, Supplemental Data File S1), we conclude that delays in the response are of minor relevance for the differences observed among the transcriptome responses.

### Identification of conserved response genes

Several gene families are known to be up-regulated by elevated auxin levels in *A. thaliana* (Paponov *et al.*, 2008). The cross-species approach might provide further insights into the identity of genes that are conserved in their response to auxin and might thus be of particular importance for auxin signaling, metabolism and/or response. The intersection of up-regulated genes among the two *Arabidopsis* species was moderate at both time points (Fig. 2C). Among the commonly up-regulated genes were individual members of prominent auxin response gene families such as the *ASYMMETRIC LEAVES/LATERAL ORGAN BOUNDARIES DOMAIN (ASL/LBD), GRETCHEN HAGEN 3 (GH3), AUX/IAA* and *SMALL AUXIN UPREGULATED (SAUR)* families (Tab. 1 and Tab. S1), validating the successful auxin induction. In addition, numerous other genes were induced by auxin treatment in both species. This included known auxin-responsive genes (e.g. *ARABIDOPSIS THALIANA HOMEOBOX 2* (*HAT2)/AT5G47370*), genes associated with other phytohormones (e.g. *1-AMINOCYCLOPROPANE-1-CARBOXYLATE SYNTHASE 11* (*ACS11)/AT4G08040, BRASSINOSTEROID INSENSITIVE LIKE 3 (BRL3)/AT3G13380, GIBBERELLIN 2-OXIDASE 8 (GA2ox8)/AT4G21200)* as well as several genes with so far unknown function (e.g. *AT1G29195, AT1G64405*, etc). The latter group might be of particular interest as the conserved response to the auxin stimulus in both species might indicate potential new candidate genes relevant for auxin responses.

**Table 1.**
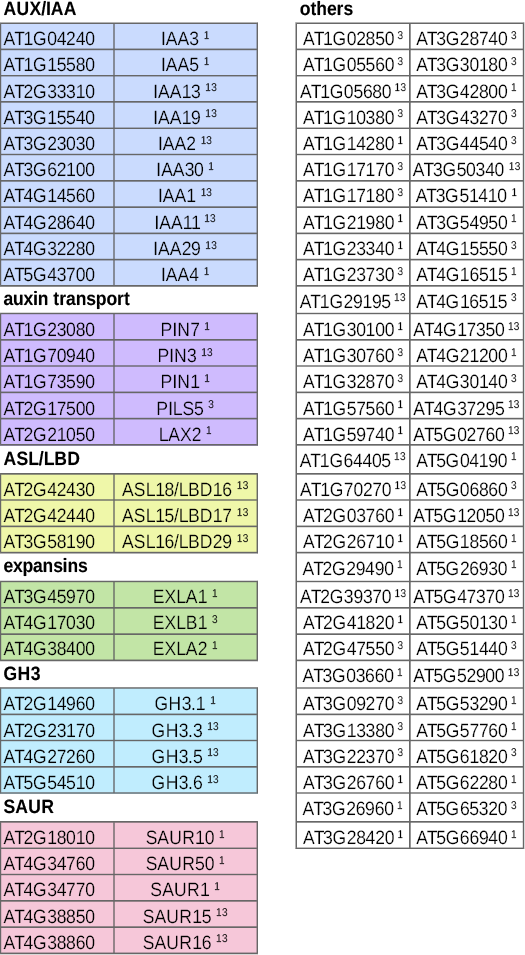
Conserved auxin up-regulated genes. Genes significantly up-regulated (log2 fold change > 1) in *A. thaliana* and *A. lyrata* after 1 h (^1^) and/or 3 h (^3^) of auxin treatment in 7 days-old seedlings. Detailed information on *A. lyrata* locus identifiers, corresponding ATH1 array elements and expression levels are shown in Tab. S1.

**AUX/IAA**
|  |  |
| --- | --- |
| AT1G04240 | IAA3<sup>1</sup> |
| AT1G15580 | IAA5<sup>1</sup> |
| AT2G33310 | IAA13<sup>13</sup> |
| AT3G15540 | IAA19<sup>13</sup> |
| AT3G23030 | IAA2<sup>13</sup> |
| AT3G62100 | IAA30<sup>1</sup> |
| AT4G14560 | IAA1<sup>13</sup> |
| AT4G28640 | IAA11<sup>13</sup> |
| AT4G32280 | IAA29<sup>13</sup> |
| AT5G43700 | IAA4<sup>1</sup> |
| **auxin transport** |  |
| AT1G23080 | PIN7<sup>1</sup> |
| AT1G70940 | PIN3<sup>13</sup> |
| AT1G73590 | PIN1<sup>1</sup> |
| AT2G17500 | PILS5<sup>3</sup> |
| AT2G21050 | LAX2<sup>1</sup> |
| **ASL/LBD** |  |
| AT2G42430 | ASL18/LBD16<sup>13</sup> |
| AT2G42440 | ASL15/LBD17<sup>13</sup> |
| AT3G58190 | ASL16/LBD29<sup>13</sup> |
| **expansins** |  |
| AT3G45970 | EXLA1<sup>1</sup> |
| AT4G17030 | EXLB1<sup>3</sup> |
| AT4G38400 | EXLA2<sup>1</sup> |
| **GH3** |  |
| AT2G14960 | GH3.1<sup>1</sup> |
| AT2G23170 | GH3.3<sup>13</sup> |
| AT4G27260 | GH3.5<sup>13</sup> |
| AT5G54510 | GH3.6<sup>13</sup> |
| **SAUR** |  |
| AT2G18010 | SAUR10<sup>1</sup> |
| AT4G34760 | SAUR50<sup>1</sup> |
| AT4G34770 | SAUR1<sup>1</sup> |
| AT4G38850 | SAUR15<sup>13</sup> |
| AT4G38860 | SAUR16<sup>13</sup> |
| **others** |  |
| AT1G02850<sup>3</sup> | AT3G28740<sup>3</sup> |
| AT1G05560<sup>3</sup> | AT3G30180<sup>3</sup> |
| AT1G05680<sup>13</sup> | AT3G42800<sup>1</sup> |
| AT1G10380<sup>3</sup> | AT3G43270<sup>3</sup> |
| AT1G14280<sup>1</sup> | AT3G44540<sup>3</sup> |
| AT1G17170<sup>3</sup> | AT3G50340<sup>13</sup> |
| AT1G17180<sup>3</sup> | AT3G51410<sup>1</sup> |
| AT1G21980<sup>1</sup> | AT3G54950<sup>1</sup> |
| AT1G23340<sup>1</sup> | AT4G15550<sup>3</sup> |
| AT1G23730<sup>3</sup> | AT4G16515<sup>1</sup> |
| AT1G29195<sup>13</sup> | AT4G16515<sup>3</sup> |
| AT1G30100<sup>1</sup> | AT4G17350<sup>13</sup> |
| AT1G30760<sup>3</sup> | AT4G21200<sup>1</sup> |
| AT1G32870<sup>3</sup> | AT4G30140<sup>3</sup> |
| AT1G57560<sup>1</sup> | AT4G37295<sup>13</sup> |
| AT1G59740<sup>1</sup> | AT5G02760<sup>13</sup> |
| AT1G64405<sup>13</sup> | AT5G04190<sup>1</sup> |
| AT1G70270<sup>13</sup> | AT5G06860<sup>3</sup> |
| AT2G03760<sup>1</sup> | AT5G12050<sup>13</sup> |
| AT2G26710<sup>1</sup> | AT5G18560<sup>1</sup> |
| AT2G29490<sup>1</sup> | AT5G26930<sup>1</sup> |
| AT2G39370<sup>13</sup> | AT5G47370<sup>13</sup> |
| AT2G41820<sup>1</sup> | AT5G50130<sup>1</sup> |
| AT2G47550<sup>3</sup> | AT5G51440<sup>3</sup> |
| AT3G03660<sup>1</sup> | AT5G52900<sup>13</sup> |
| AT3G09270<sup>3</sup> | AT5G53290<sup>1</sup> |
| AT3G13380<sup>3</sup> | AT5G57760<sup>1</sup> |
| AT3G22370<sup>3</sup> | AT5G61820<sup>3</sup> |
| AT3G26760<sup>1</sup> | AT5G62280<sup>1</sup> |
| AT3G26960<sup>1</sup> | AT5G65320<sup>3</sup> |
| AT3G28420<sup>1</sup> | AT5G66940<sup>1</sup> |

### Inter-species expression responses in auxin-relevant gene families

To further investigate similarities and specificities of transcriptional auxin responses in *A. thaliana* and *A. lyrata*, we performed a cluster analysis of genes that showed a change in expression in at least one species at any of the analyzed time points with a coefficient of variation (cv) > 0.05. A modified Pearson correlation (*mod.r*) was used as a distance measure in the hierarchical clustering to incorporate information on the variation among the three biological replicates at each analyzed time point. To filter for correlations among genes with potential biological relevance, we further applied a minimum cut-off in correlation of *mod.r* = 0.7. The resulting 14 gene clusters fall into two clearly distinct groups (Fig. 3). Clusters 1 - 8 and clusters 9 - 14 are predominantly characterized by genes that show a higher expression level and/or response in *A. lyrata* or *A. thaliana*, respectively. Only very few clusters show high similarities among the expression profiles of both species (e.g. cluster 2 and 9). The majority of cluster profiles show small to striking differences between the two species in either expression levels (e.g. cluster 8) or expression response in terms of induction/repression profiles (e.g. cluster 3) or both (e.g. cluster 11). We next inspected whether the presence and frequency of known *cis*-regulatory elements in the promoters of clustered genes could explain the observed patterns of similarities or differences in the expression profiles of individual clusters. We limited the size of the putative promoter region to 500 bp upstream of the transcription start site. While eukaryotic promoters can arguably be much larger, the majority of *cis*-regulatory sequences should be present within this 500 bp interval (Franco-Zorrilla *et al.*, 2014). We analyzed the presence of 99 known *cis*-regulatory elements taken from the *Arabidopsis cis*-regulatory element database (http://arabidopsis.med.ohio-state.edu/AtcisDB/) and additional literature (Tab. S2). Of the total number of motifs (n = 109) 35 known *cis*-elements were detected in at least one of the promoter sequences of clustered genes with significantly altered expression (Tab. S3). To assess whether the presence of certain regulatory sequences explains the distinct expression profiles, we initially focused on *cis*-elements known or predicted to be involved in auxin responses such as different varieties of the auxin responsive element (AuxRE), the E-box/hormone up at dawn (HUD) element and the TGA2 binding site motif (Liu *et al.*, 1994; Nemhauser *et al.*, 2004; Vert *et al.*, 2008; Keilwagen *et al.*, 2011).

**Fig. 3.**
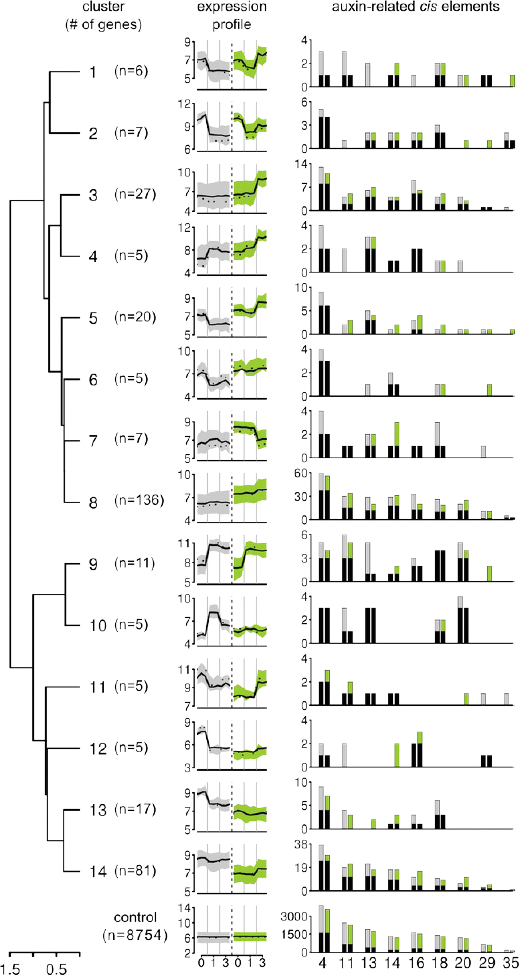
Cluster analysis of auxin-regulated genes and allocation of known *cis*-regulatory elements. Hierarchical clustering of genes that showed an auxin-induced expression response (coefficient of variation (cv) > 0.05) in at least one species at one time point of auxin treatment using a modified Pearson correlation (1-*mod.r*) among expression profiles as distance measure. A threshold of 1-*mod.r* = 0.3 provided 14 clusters. Expression profiles show mean (solid lines) and median (dotted lines) expression levels of genes in one cluster. Areas shaded in gray and green denote interquartile ranges for *A. thaliana* and *A. lyrata*, respectively. Bar plots illustrate the presence of known *cis*-element sequences with functional relevance in auxin biology. “4”: AATAAG, “11”: TGTCTC, “13”: CACATG, “14”: CGTG[TC]G, “16”: CACCAT, “18”: TGTCTG, “20”: TGT[CG]T[CG][CGT]C, “29”: TGTATATAT, and “35”: ATACGTGT. A full description of *cis*-elements is shown in Tab. S2 and S3. A comprehensive analysis of the presence of known regulatory sequences is depicted in Fig. S3.

Auxin-related *cis*-regulatory elements were detected in all of the clusters. There was a certain degree of redundancy in the analysis due to sequence overlaps among different variants of elements, e.g., in various versions of the AuxRE (Nos. 11, 18, and 20; Fig. 3, Tab. S3). Yet, neither the frequency of AuxREs nor any other *cis*-element seemed to explain the similarities or differences in the expression behavior (i.e., auxin response pattern) of the gene clusters (Fig. 3, Fig. S3). Even for cluster 9, which shows clearly up-regulated profiles in both species and includes several prominent auxin-responsive genes, only roughly 50% of the genes contained a version of the AuxREs. This observation is in accordance with several previous studies in *A. thaliana* which showed a lack of AuxREs in a substantial number of auxin-regulated genes (Nemhauser *et al.*, 2004). Furthermore, expression differences among *A. thaliana* and *A. lyrata* did not show a clear pattern of correlation to the species-specific presence of individual regulatory elements in the promoters of *A. thaliana* (gray) or *A. lyrata* (green). However, these observations remain subjective as statistical tests for over- or under-representation of elements are hindered by the low number of genes present in several of the clusters identified here.

### Expression divergence vs. promoter divergence

The lack of any obvious correlation of known *cis*-elements and auxin-induced expression patterns prompted a *de novo* search for putative regulatory sequences. The data set seemed ideal as the two *Arabidopsis* species are distant enough to provide considerable sequence variation in promoter regions while providing sufficient similarities to allow for local alignments of the sequences (Hu *et al.*, 2011). However, a prerequisite for this approach would be a general correlation between the diversity in the promoter sequence and the differences detected on the expression level. To evaluate this assumption, we compared promoters of three groups of genes: (i) the set of conserved genes with a significant induction in expression in response to 1 h of auxin treatment in both species (n = 68), (ii) promoters of genes that are up-regulated in at least one of the analyzed species (n = 297), which include also the 68 genes of group (I) that met the threshold of auxin-induction in both species. We retained this gene set in group (ii) as the kinetics of expression profiles might still show differences among the two species. Group (iii) included neutral genes that did not show a significant alteration in expression as a control set (n = 11195). We then calculated the expression divergence of expression profiles between each orthologous gene pair using *mod.r*. Similarities of promoter sequences were assessed by a sliding window approach to compute the correlation of the occurrence of all possible 8-mers across the promoters of orthologous genes (*kmer.r,* Vinga and Almeida, 2003).

As expected, expression divergence for genes with a conserved up-regulation in both species is rather low and seems to be independent of promoter divergences (Fig. 4A). Similarly, no correlation among expression and promoter divergence was observed for neutral genes that did not show expression changes in response to auxin. However, for group (ii) including all genes with a differential response in at least one of the two analyzed species, a wide range in expression divergence as well as promoter divergence was observed which showed a considerably higher correlation compared to the other two gene sets (Fig. 4A). Hence, both auxin-responsive gene sets showed the expected pattern of relationships between expression and promoter divergence, which made them suitable candidate sets for *de novo* identification of regulatory promoter elements.

**Fig. 4.**
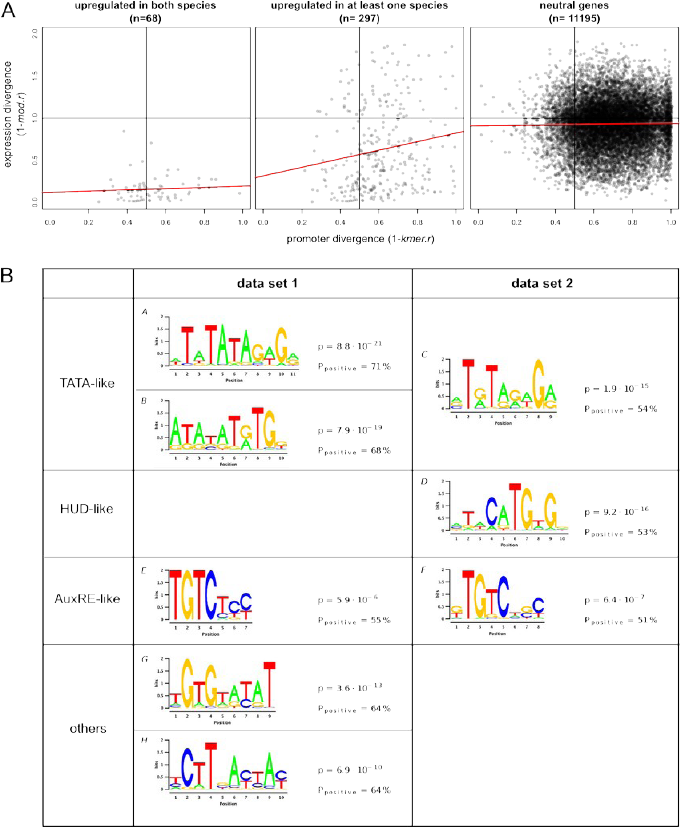
*De novo* identification of promoter elements. (A) Analysis of promoter and expression diversity in genes that are significantly up-regulated in both species, up-regulated in either *A. thaliana* or *A. lyrata* or non-responsive (neutral) to 1 hour of auxin treatment. Divergence among expression profiles and promoter sequences was assessed by *mod.r* correlation of expression profiles and 8-mer sliding window correlation (*kmer.r*) results of promoter sequences, respectively. (B) *De novo* identification of putative *cis*-regulatory elements that are significantly overrepresented in auxin-induced genes using Dimont. Motifs shown were significantly enriched in genes up-regulated in both species (data set 1) or in at least one species (data set 2). Motifs were additionally tested for enrichment in an independent auxin-induced expression data set of *A. thaliana* (see *p’* values in Fig. S4). Frequency of occurrence [%] in the positive and control data sets are denoted by *%_positive_* and *%_control_*, respectively.

### De novo identification of putative cis-regulatory elements

Based on the promoter divergence analysis we selected two gene sets for motif discovery (positive data sets). The first set comprised an extended set of genes that were induced in both species after 1 h of auxin treatment. As we did not limit the selection by filtering via corrected p-values, this set extended the previously shown set of genes of up-regulated in both species to a total of 81 orthologous gene pairs. Data set 2 comprises promoters of an extended set of genes that were up-regulated in at least one species. For this second data set we only included the promoter sequence of the species that showed a significant up-regulation of a gene in response to auxin (n = 845 promoter sequences). The corresponding promoter sequence of the other species of an orthologous gene pair was included in the control data set 2 following the rationale that regulatory elements required for the auxin response are absent in this case.

Applying the discriminative motif discovery tool *Dimont* (Grau *et al.*, 2013), we identified motifs with significant over-representation in each of the two data sets of auxin-induced genes in comparison to their respective control data sets (see Methods for details). Among the motifs identified in both data sets were sequences with high to medium similarities to TATA box elements (Fig. 4B, motifs A - C). TATA boxes are present in approximately 28 % of all *Arabidopsis* genes with a predominance of non-housekeeping genes (Molina and Grotewold, 2005). Interestingly, yeast genes containing a TATA box showed increased inter-species variation in expression responses to a variety of environmental stresses (Tirosh *et al.*, 2006). It was hypothesized that core promoters including a TATA box might be more sensitive to genetic perturbations and could be a driving factor in expression divergence (Tirosh *et al.*, 2006). As TATA-like elements were enriched in both analyzed data sets they might rather reflect the general rapid and partially strong induction of these genes in response to an external stimulus. In yeast, TATA-containing promoters showed a slightly higher tendency for higher expression after a heat shock (Kim and Iyer, 2004). The identification of novel variants of AuxRE-and HUD-like motifs (Fig. 4B, motifs D - F) corresponds with their previously demonstrated function in auxin-mediated expression induction (Walcher and Nemhauser, 2012) and indicates a general success in the analytical approach. The identification of these putatively novel variations of known elements may indicate a higher tolerance for sequence variation in the *cis*-regulatory motif that only becomes evident with a higher degree of genetic variation among genome sequences included in this analysis. Recent advances in understanding the mode of ARF transcription factor binding to target promoter sequences substantiates this assumption. Structure-function analysis indicated that different ARF proteins seem to have altered affinities for different variations of AuxREs (Boer *et al.*, 2014). These specificities could account at least partially for functional specifications of individual ARFs and might also be a contributing factor in natural variation of transcriptional auxin responses.

In addition, other putatively novel *cis*-regulatory sequences were found to be significantly enriched in genes that were induced by auxin in both species (Fig. 4 and Fig. S4, motifs G - L). To the best of our knowledge, these sequences have not been described previously. To assess the potential significance of these elements with respect to auxin responses, we tested whether they were also enriched in auxin-induced genes in an independent auxin response transcriptome dataset generated for *A. thaliana* seedlings (Nemhauser *et al.*, 2006). Two of the sequences (Fig. 4B, motifs G + H) were indeed found to be enriched significantly (p < 0.05) in differentially expressed genes in this additional data set (Fig. S4), highlighting their potential relevance for auxin-induced transcriptional regulation. We then inspected whether the presence/absence of any of the *de novo*-identified promoter sequences can account for the differential expression responses or levels of distinct gene clusters (Fig. S3). However, similarly to the analysis of previously described *cis*-elements, no coincidence pattern of *de novo* promoter elements and expression response could be identified despite the enrichment of these sequences in auxin-regulated genes. While we cannot exclude that the newly identified promoter sequences may be of minor functional relevance, the analysis as a whole rather points towards a highly complex orchestration of auxin-induced expression responses involving multiple *cis*-element variations.

The diversity in auxin-induced expression responses via combinations of multiple different transcription factors and their individual target promoter sequences has been shown previously in case of the AuxRE and HUD elements which seem to be acting interdependently in facilitating efficient ARF binding (Walcher and Nemhauser, 2012). G-box and Myb-binding motifs have similarly been described to function as components in composite AuREs (Ulmasov *et al.*, 1995) and ABRE-like and Y-patch motifs have recently been identified in a bioinformatics approach as putative constituents of composite AuxREs (Mironova *et al.*, 2014). Composite AuxREs may be integral to form the auxin code and it is possible that several as yet uncharacterized motifs contribute to the diversity. Furthermore, different types of AuxREs seem to contribute differentially to transcriptional auxin responses (Zemlyanskaya *et al.*, 2016), putatively via preferential binding of inducing or repressing ARFs. While the well known canonical AuxRE TGTCTC is found in the regulatory regions of up-regulated genes of the early auxin response (0.5 - 2 h), the motif TGTCAT is found in down-regulated genes of the late (4 - 24 h) auxin response. The motif TGTCGG is not as strongly restricted to specific conditions as it is found in up-regulated genes of both the early and late auxin response (Zemlyanskaya *et al.*, 2016).

Unraveling the combinatorial code of regulatory elements will require highly sophisticated bioinformatic approaches, a higher number of transcription profile data sets from diverse genetic backgrounds and preferably from distinct tissue sets for in-depth phylogenetic footprinting analyses and ultimately extensive functional validation.

While the complex promoter code of auxin-induced transcriptional variation remains somewhat elusive, the general hierarchy of the auxin signal transduction pathway is well known. Transcriptional responses to auxin are primarily mediated via the TIR1/AFB-AUX/IAA-ARF signaling pathway. All three components are encoded by gene families. Individual members of these families seem to have partial redundancies in their spatio-temporal expression patterns and have at least partially distinct biochemical properties (Okushima *et al.*, 2005; Paponov *et al.*, 2008; Parry *et al.*, 2009; Rademacher *et al.*, 2011; Calderón Villalobos *et al.*, 2012). As quantitative alterations in the equilibrium of these signaling components may significantly affect downstream responses, we next focused on this particular group of genes.

### Divergence of AUX/IAA gene expression is reflected in downstream responses

Diversity in co-expression profiles of signaling components have previously been shown among different accessions of *A. thaliana* (Delker *et al.*, 2010). Variation in gene expression levels and co-expression patterns are indicative of altered levels of individual signaling proteins that might contribute to the differential responses observed initially on transcriptional and ultimately on physiological levels (Delker *et al.*, 2010). Differential expression was predominantly evident for *AUX/IAA* genes which are generally more responsive to auxin treatment than *ARFs* or *TIR1/AFBs* (Paponov *et al.*, 2008). Variation of *AUX/IAA* transcriptional activation is indicative of differential signal transduction events in response to a similar stimulus. *AUX/IAA* genes constitute primary auxin response genes that provide a read-out for the activation of the auxin signal transduction pathway. Subsequent alterations in AUX/IAA protein levels will likely impact further on auxin sensing by affecting the availability of individual auxin co-receptor complexes with potentially specific auxin sensitivities. Preferential formation of specific ARF-AUX/IAA heteromerizations may additionally affect transcriptional regulation. As such, the intra-specific comparison of auxin-regulated expression responses in *A. thaliana* accessions highlighted the early auxin signaling network as a potential source for the observed variation in downstream responses (Delker *et al.*, 2010). In this study, we challenged this hypothesis by inspecting the expression responses of the core auxin signaling gene families in the cross-species comparison of auxin responses.

Members of all three gene families showed differential expression responses between the two species. Analysis of expression and promoter divergences showed a considerably stronger correlation for the highly auxin-responsive *AUX/IAA* gene family (Fig. 5A). This might be similar for the *TIR/AFB* family but the total number of only four genes retained in this analysis is too low and effects by individual outliers may be high. While promoter divergences of *ARF* family members are also quite high, expression divergence is only low to medium (1-*mod.r* values in expression divergence from 0-1, Fig. 5A). AUX/IAAs have a unique role among the signaling components. Apart from their dual function in signaling as repressors of ARF transcription factors and co-receptors of auxin, they also constitute a group of classic and conserved auxin response genes which provide a readout for auxin responsiveness (Tab. 1, Paponov *et al.*, 2008). Due to this prominent role, we inspected the expression responses of the *AUX/IAA* gene family in more detail. Hierarchical clustering allowed the identification of *AUX/IAA* subgroups based on the correlation (1-*mod.r*) in expression profiles (Fig. 5B). While clusters 1 - 3 contained *AUX/IAA* genes that were induced by auxin only in *A. lyrata*, clusters 5 - 8 *AUX/IAA* genes responded primarily in *A. thaliana.* In contrast, cluster 4 contained *AUX/IAA* genes that showed significantly changed expression levels in response to auxin treatment in both species. These genes are part of the conserved auxin response gene set (Tab. 1) and form the largest cluster among the *AUX/IAA* genes (Fig. 5B). Consequently, *AUX/IAA* genes with similar expression profiles in *A. thaliana* and *A. lyrata* are indicative for similar upstream transcriptional activation/signaling events and their corresponding gene products can be speculated to have similar downstream signaling effects. In contrast to that, gene clusters with species-specific auxin responses could be indicative for the sources of natural variation seen in downstream auxin responses.

**Fig. 5.**
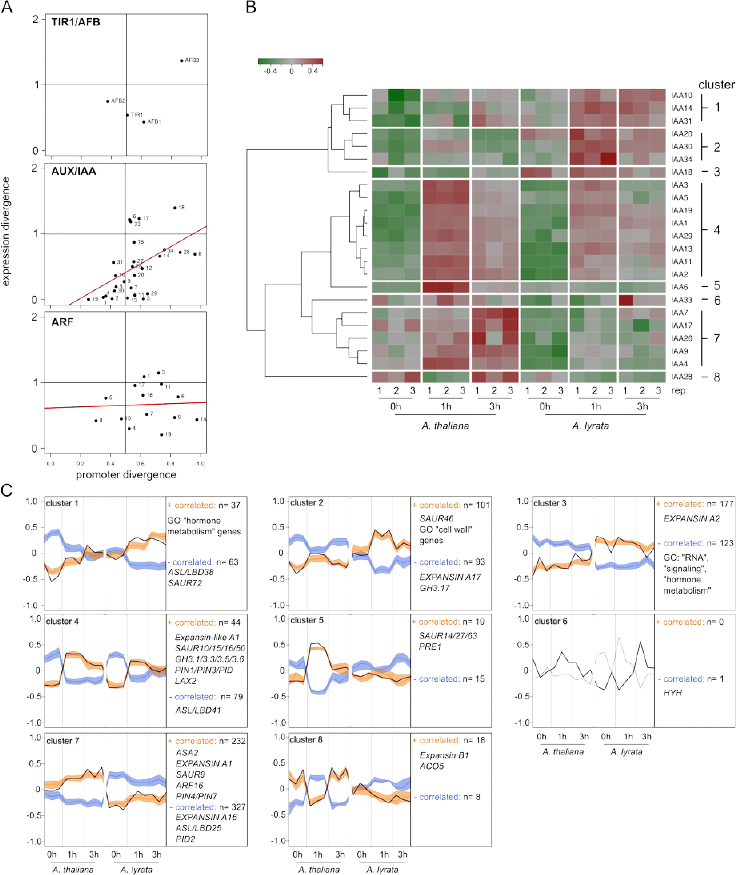
*AUX/IAA* expression divergence correlates with downstream expression profiles. (A) Promoter divergence for core auxin signaling genes was determined as described in Fig. 4A. (B) Hierarchical clustering of PIF-normalized (mean-centered) *AUX/IAA* expression profiles using 1-*mod.r* as distance measure. (C) Selected genes with expression profiles that are positively (+) or negatively (-) correlated to the mean expression profiles (solid black lines) of *AUX/IAA* clusters shown in B as determined by the profile interaction finder (PIF) algorithm. A complete list of identified genes is presented as Data File S2.

To identify genes with expression profiles that are either positively or negatively correlated to individual *AUX/IAA* gene clusters (Data file S2), we used the recently introduced *Profile Interaction Finder* (*PIF*) algorithm (Poeschl *et al.*, 2014). As expected, members of several of the classic and conserved auxin response gene families showed positively correlated expression profiles to cluster 4 (Fig. 5C). This cluster shows a classic response profile of transient expression induction in both species. The respective *AUX/IAA* and co-regulated genes of known auxin-related genes seem to be part of a conserved auxin response in both species.

Clusters with more species-specific expression responses also showed correlations with genes relevant for auxin biology. For example, the expression profile of cluster 7 shows a higher expression and gradual auxin induction in *A. thaliana,* while the expression levels in *A. lyrata* are generally lower. A similar, positively correlated pattern in expression was observed for several auxin-relevant genes ranging from biosynthesis (*ASA2*), to signaling (*ARF16*), transport (*PIN4, PIN7*), and response (*EXPANSIN A1*). In addition, genes with negatively correlated expression profiles were also identified (e.g. *ASL/LBD25*).

To assess the wider implications of *AUX/IAA* expression variation for downstream response patterns, we compared the inter-species variation of this data set with the intra-species variation among seven *A. thaliana* accessions of a previous analysis (Delker *et al.*, 2010). Expression divergence of *AUX/IAA* genes revealed a group of *AUX/IAAs* with highly conserved expression responses corresponding primarily to genes sorted to cluster 4 (Fig. 5B, Fig. 6A). Other genes showed higher and similar inter- and intra-species divergence (e.g. *AUX/IAA20* and *34*). Finally, a group of 5 *AUX/IAA* genes showed a higher diversity in the comparison among *A. thaliana* and *A. lyrata* than in the intra-species comparison. These genes cause a considerable increase in the inter-species divergence on *AUX/IAA* level (Fig. 6A).

**Fig. 6.**
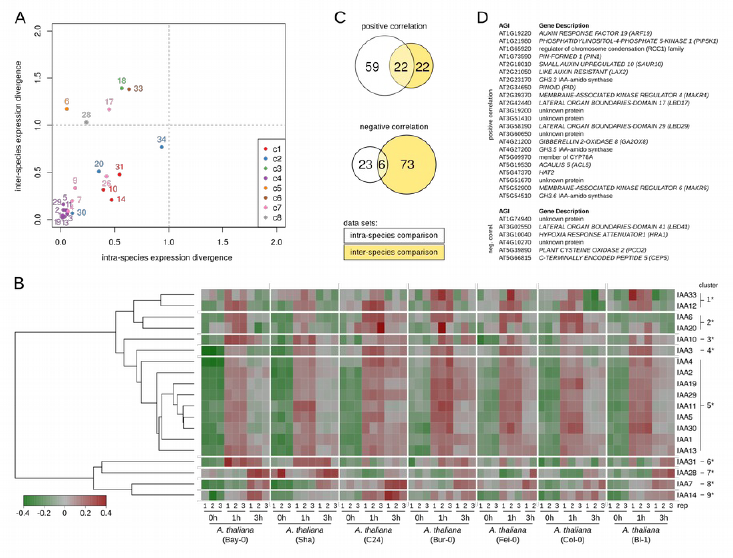
Inter- and intra-species expression divergence of *AUX/IAA* genes. (A) Comparison of mean *AUX/IAA* expression divergence detected in the inter-species comparison of *A. thaliana* vs. *A. lyrata* and an intra-species comparison among seven *A. thaliana* accessions. *AUX/IAAs* are color-coded according to their cluster number in Figure 5B. (B) Independent hierarchical clustering of PIF-normalized (mean-centered) intra-species *AUX/IAA* expression profiles using 1-*mod.r* as distance measure. Asterisks mark cluster numbers of the intra-species data set to allow differentiation from clusters in Figure 5B. (C) Venn diagrams illustrate the overlap of genes with expression profiles that are positively (+) or negatively (-) correlated to the mean expression profiles of *AUX/IAA* clusters 4 (Figure 5B) and cluster 5*. (D) List of gene identities with overlapping correlations. A complete list of specific and overlapping genes is shown in Supplemental Data File 2.

To account for potential differences in cluster structures, *AUX/IAA* genes were independently clustered based on their expression behavior in the seven *A. thaliana* accessions (Fig. 6B). As one of the prerequisites in gene selection was a cv > 0.05, only 19 *AUX/IAA* genes remained in the intra-specific cluster analysis whereas *AUX/IAA 9, 18* and *26* did not show auxin-induced changes in expression profiles. We marked the new cluster with an “*” to differentiate between the two cluster analyses. The largest cluster (cluster 5*) again is formed by *AUX/IAAs* showing a uniform transient induction of expression. This cluster largely corresponds to genes grouped in cluster 4 of the inter-species comparison but includes additional genes (*e.g., AUX/IAA 30*) which clustered differently in the other data set. Apart from the conserved cluster, other *AUX/IAAs* show differential expression patterns in different accessions and seem to be highly variable in their response which may contribute to differential downstream responses (Fig. 6B). To assess potential implications on downstream responses, we searched for positively and negatively correlated genes using the *PIF* algorithm. We focused on cluster 5* here, as the constitution and expression profiles of the genes were comparable to cluster 4 in the inter-species comparison (Fig. 5B, Fig. 6B). Similar to the previous analysis, a number of known auxin-responsive genes were identified to be either positively or negatively correlated to cluster 5* (Supplemental Data set 2). Comparing the lists of cluster 4 and cluster 5* showed a high overlap in the positively correlated genes (Fig. 6C) and a much lower overlap in negatively correlated genes. Yet, both lists contain known auxin-responsive genes and even more interesting, genes which have not been characterized or linked to auxin so far (Fig. 6D). The fact that these genes show a reproducible strong correlation with classic auxin response genes may indicate a function in auxin biology that awaits unraveling.

The correlation of numerous auxin-associated genes with *AUX/IAA* gene clusters indicates that variation in early auxin signaling may penetrate to downstream response levels. Ultimately, these differences could quantitatively contribute to the variation observed on physiological levels. Whether the major source of variation is actually caused by differential expression or rather by altered biochemical properties due to non-synonymous mutations of signaling genes remains to be elucidated. The genome-wide variation in auxin-induced gene expression may originate in the differential gene regulation and subsequent protein levels of AUX/IAAs themselves. Alternatively and/or in addition, differential upstream events such as auxin sensing or initial gene activation by ARFs may be the actual source of initial variation which then results in differential activation of *AUX/IAAs* and other genes.

## Summary and conclusions

We studied inter-species variation of physiological and transcriptional auxin responses to assess whether the highly conserved auxin signaling and response pathway might contribute to adaptive processes in growth and development. Transcriptome analysis allowed the identification of genes with a highly conserved response to the auxin treatment which included both, members of known auxin-responsive gene families and so far uncharacterized genes. However, the majority of differentially expressed genes in response to auxin showed significant variation in expression levels and/or response patterns between the two *Arabidopsis* species. Neither similar nor species-specific expression patterns of auxin-regulated gene clusters could be explained by the presence of individual known or *de novo*-identified promoter elements. Thus, it remains likely that a complex code of element combinations accounts for the diversity in transcriptional auxin responses. Breaking this particular code will require extensive efforts by bioinformaticians and far more available expression data from genetically diverse backgrounds.

A significant source for variation in auxin-induced transcriptome changes likely originates within the initial auxin signal transduction pathway itself. Distinct patterns of *AUX/IAA* gene cluster expressions were found to penetrate to the level of numerous response genes, many of which with a known functional relevance for auxin biology. While *AUX/IAA* gene expression divergence may contribute directly to differential activation of downstream responses, it is also indicative for species-specific differences by which identical auxin signals are transduced into gene expression responses. Consequently, the triumvirate of TIR1/AFBs, AUX/IAAs, and ARFs harbor significant potential for the initiation of variation in downstream auxin signaling and response.

## Supplemental Material

**Supplemental Methods:** Comprehensive description of *de novo* identification of *cis*-elements

**Supplemental Tab. S1:** Expression response of conserved auxin up-regulated genes in *A. thaliana* and *A. lyrata*

**Supplemental Tab. S2:** collection of known *cis*-regulatory elements

**Supplemental Tab. S3:** *cis*-regulatory elements identified in significantly up-regulated genes

**Supplemental Data File S1:** Tables of common and species-specific up-regulated genes including cross-comparisons

**Supplemental Data File S2**: Profile interaction finder results for genes positively or negatively correlated with *AUX/IAA* gene clusters

**Supplemental Fig. S1:** Absolute lengths in physiological auxin responses, growth dynamics, auxin uptake and cross-comparison of up-regulated genes

**Supplemental Fig. S2:** Expression levels of non-responsive reference genes confirm successful normalization of the cross-species microarray data

**Supplemental Fig. S3:** Assignment of 35 known and 8 *de novo*-identified *cis*-elements to auxin-regulated gene clusters

**Supplemental Fig. S4:** *De novo-*identified *cis*-elements

## Acknowledgements

This work was supported by the Deutsche Forschungsgemeinschaft (Qu 141/3-1 to MQ) and by core funding of the Leibniz association. We furthermore thank Ivo Grosse for fruitful discussions in the initial project phase.

